# An ancient role for the Hippo pathway in axis formation and morphogenesis

**DOI:** 10.1101/2022.01.19.476962

**Authors:** Maria Brooun, Willi Salvenmoser, Catherine Dana, Marius Sudol, Robert Steele, Bert Hobmayer, Helen McNeill

## Abstract

How did cells of early metazoan organisms first organize themselves to form a body axis? The canonical Wnt pathway has been shown to be sufficient for induction of axis in Cnidaria, a sister group to Bilateria, and is important in bilaterian axis formation. Here, we provide experimental evidence that in cnidarian *Hydra* the Hippo pathway regulates the formation of a new axis during budding upstream of the Wnt pathway. The target of Hippo pathway, the transcriptional co-activator YAP, inhibits the initiation of budding in *Hydra*, and is regulated by *Hydra* LATS. In addition, we show functions of Hippo pathway in regulation of actin organization and cell proliferation in *Hydra*. We hypothesize that Hippo pathway served as a link between continuous cell division, cell density and axis formation early in metazoan evolution.

## Introduction

How animals establish and pattern their primary body axis is a fundamental problem in biology. Data from a wide range of bilaterian animals suggest that Wnt signaling controls posterior identity during body plan formation [1-3]. Wnt also drives axis formation in Cnidaria, the sister phylum to Bilateria. [4-7]. Thus, an axial patterning role for Wnt signaling was present in the last common ancestor of Cnidaria and Bilateria, which diverged ~ 650 million years ago.

*Hydra*, a small freshwater cnidarian with a simple body plan, exhibits amazing regenerative and budding capabilities, as described in 1744 by Trembley [8]. The *Hydra* body is essentially a tube of epithelial cells, aligned along the oral-aboral axis (Figures 1A and B). *Hydra* has only two epithelial layers, an ectoderm and endoderm, separated by an extracellular matrix called the mesoglea (Figure 1C). Epithelial cells of both layers are attached to the mesoglea and exhibit actin-myosin contractile elements called muscle processes on their basal sides (Figures 1C and D). Cells of the interstitial cell lineage, stem cells which give rise to nerves, nematocytes, gland cells, and germ cells cells [9], are intermingled among the epithelial cells (Figure 1C). Epithelial cells continuously divide and are displaced along the oral-aboral axis towards the head and foot. The balance between cell production and loss of cells via sloughing from the ends and bud formation determines the size of the animal [10, 11] (Figure 1B).

**Figure 1.**
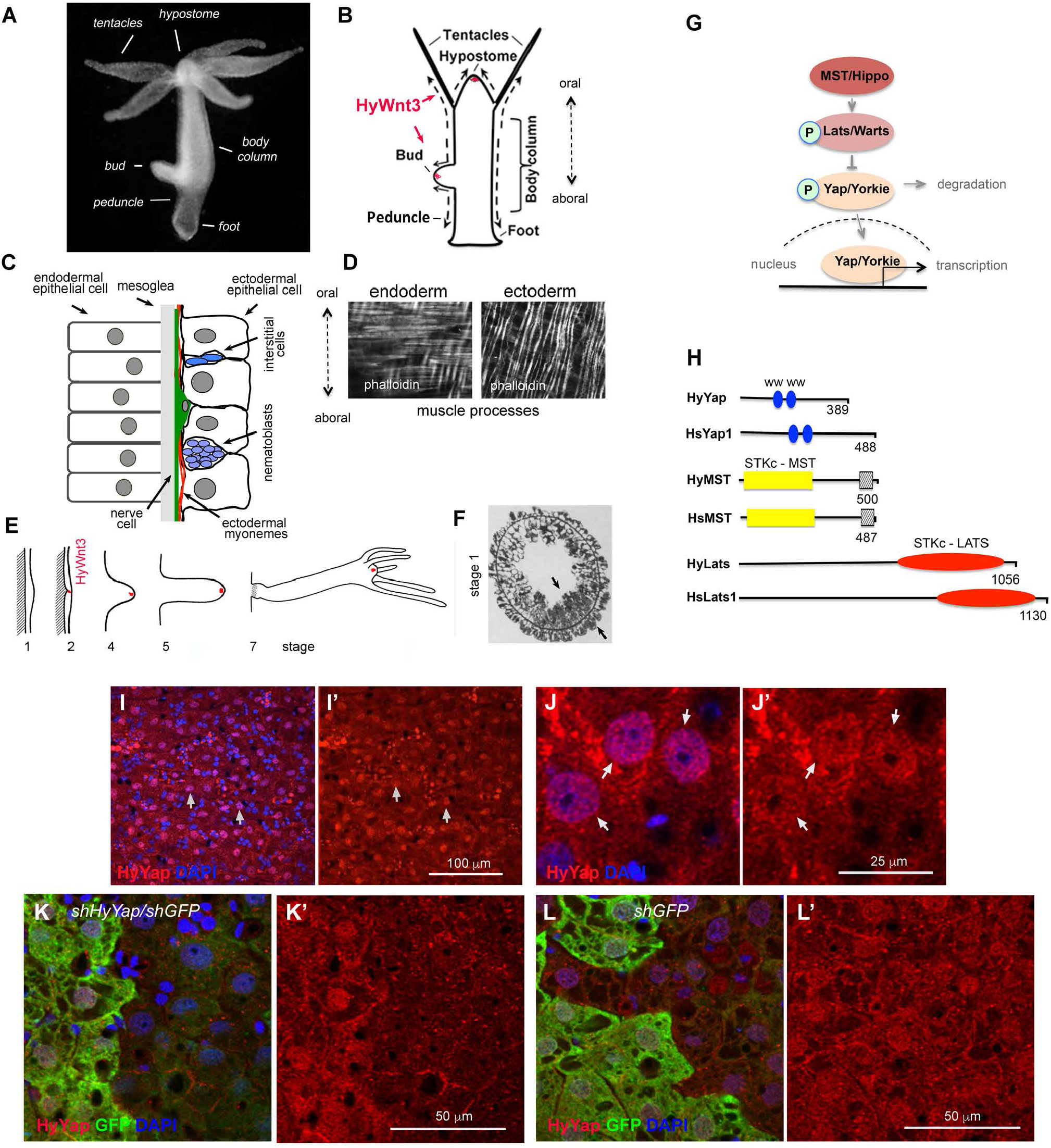
Hydra homologue of YAP is expressed in ectodermal epithelial cells. (A) Photo of a live *Hydra*. (B) Schematic of *Hydra* body plan; arrows indicate directions of cell displacement along the oral/aboral axis. (C) Schematic of a section through *Hydra* body column. (D) *Hydra* endodermal and ectodermal muscle processes visualized with phalloidin. (E) Schematic of *Hydra* budding (adapted from [19]). The area of expression of *HyWnt3* is confined to about 50 ectodermal epithelial cells marked in red [14]. (F) Stage 1: transverse section through the budding zone, arrows indicate increased cell density in the ectoderm and endoderm (adapted from [20]). (G) Schematic of Hippo pathway. (H) Schematic of *Hydra* and mammalian homologues of Yap, MST and LATS proteins; WW – proline-rich sequences binding domain, STKc–MST1/2 catalytic domain of MST family of serine/threonine kinases,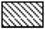-MST1-SARAH – apoptosis-mediating domain, STKc–LATS – catalytic domain of LATS family of serine/threonine kinases. (I – J’) Apical view of *Hydra* ectoderm immunostained with anti-HyYap serum. Arrows point to the nuclei of ectodermal epithelial cells. (K - L’) Apical view of ectoderm of *GFP* polyp electroporated with *shGFP/HyYap* (K, K’), *shGFP* alone(L, L’) hairpins and immunostained with anti-GFP and anti-HyYap antibodies.

Tentacles and buds form by tissue evagination. Budding, *Hydra*’s asexual form of reproduction, generates a new new body axis [12] (Figures 1B and E). Bud induction depends on the size of the mother polyp and its epithelial growth rate [13]. Budding becomes visible by a thickening of the ectoderm at a site in the lower half of the body column (stage 1 in Figure 1E; Figure 1F) and continues as evagination of both layers. The formation of a new axis in *Hydra* is also controlled by the Wnt/β-catenin pathway [4, 6]. Experimental activation of Wnt signaling in the body column results in formation of ectopic axes [4, 6, 14-16]. Expression of *HyWnt3* is detected early in budding, but only after thickening of the ectoderm is visible [14, 17]. It is unknown what pathway(s) leads to the thickening of the ectoderm and induction of *HyWnt3*.

The Hippo pathway is a key regulator of cell proliferation, differentiation, and apoptosis [18]. It consists of a cascade of kinases that controls nuclear localization of the transcription factor Yap (Yorkie in *Drosophila*) (Figure 1G). MST (*Drosophila* Hippo) kinase phosphorylates and activates LATS (*Drosophila* Warts) that, in turn, phosphorylates Yap (Yorkie) leading to its binding to 14-3-3 and cytoplasmic retention or ubiquitination and degradation [21]. Nuclear Yap/Yorkie, which promotes the expression of pro-proliferative and anti-apoptotic genes, is involved in regulation of organ size [21, 22] and morphogenesis [23].

The Hippo pathway is remarkably conserved in metazoans and their unicellular ancestors [24]. Functional studies using Hippo, Warts, and Yorkie homologues from the unicellular holozoan *Capsaspora owczarzaki* proteins ectopically expressed in *Drosophila* indicate the growth-regulating capabilities of the Hippo pathway were established before the emergence of metazoans [24]. The full repertoire of proteins making up the Hippo cascade has been identified in Ctenophora and Cnidaria [25, 26]. Immunostaining analysis of Yap in the cnidarian *Clytia hemisphaerica* suggested that regulation of nuclear/cytoplasmic localization of Yap could be a mechanism halting cell division and triggering differentiation programs [25]. However, the function of the Hippo pathway has not been elucidated in cnidarians.

Here, we investigate the role of the Hippo pathway in morphogenesis and axis formation in *Hydra*. Using *shRNA*-mediated knockdown as well as a gain of function transgenic approach, we show that the Hippo pathway components LATS and YAP regulate axis formation and morphogenesis in *Hydra*. Our studies indicate that the Hippo pathway affects epithelial growth and acts upstream of *Wnt3a* during axis formation in *Hydra*, suggesting that linkage between these two signalling pathways was a key step in the evolution of axis formation in metazoans.

## Results

### *HyYap* regulates proliferation in *Hydra*

Transcriptomic and genomic data identified single homologues of Yap, Lats and MST in *Hydra vulgaris*, which we refer to as HyYap, HyLATS and HyMST (Figure 1H and Figues S1A, B and C), consistent with recent studies [27]. To explore HyYap function *in vivo*, we generated antiserum against residues 1 – 159 of the protein. Immunostaining of *Hydra* polyps revealed HyYap in the nuclei and cytoplasm of ectodermal epithelial cells (Figures 1I - J’). Both nuclear and cytoplasmic staining were specifically removed by pre-absorption of anti-HyYap serum with HyYap-GST antigen (Figure S1E). The distribution of HyYap was similar to the subcellular distribution of mammalian and *Drosophila* Yap homologues [28, 29]. Immunoblotting of *Hydra* lysates with the anti-HyYap serum revealed a single strong band which was lost by pre-absorption with HyYap-GST antigen (Figure S1D) and by *shRNA* knockdown (see below).

To explore the function of *HyYap*, we combined *shRNA* knockdown protocols developed for *Hydra* [30, 31] and for the sea anemone *Nematostella vectensis* [32]. To optimize the protocol, we used a transgenic *Hydra* line expressing GFP in the ectodermal epithelial cells [33]. Transgenic polyps were electroporated with *shGFP*, leading to mosaic downregulation of GFP. Downregulation of GFP was seen on one side of 60 – 70% of the electroporated polyps 4 – 5 days after electroporation, and was visible 4 weeks after electroporation (Figure S2A). These data indicated that electroporation is an effective way to generate mosaic loss of function in *Hydra*. Mosaic loss of function has been a powerful tool in *Drosophila*, and we show here that this is also the case in *Hydra*.

To knock down *HyYap*, GFP-expressing polyps were electroporated with a mixture of *shGFP* hairpins and two different *shHyYap* hairpins, *shHyYap1* and *sHyhYap2* (Table S1, *shHyYap* further in text). Electroporation with a combination of *shGFP and shHyYap* dramatically reduced both nuclear and cytoplasmic HyYap staining (Figures 1K and K’). Importantly, HyYap staining was not affected by electroporation with a combination of *shGFP and shHyYapscr* or *shGFP* alone (Figures 1L, L’ and S2B). Knocking down of *HyYap* was confirmed by qPCR and immunoblot analyses (Figures S2C - E). Importantly, all *GFP*-negative cells were also *HyYap*-negative, i.e. all affected cells received both *shGFP* and *shHyYap* hairpins, leading to a reduction in the level of both proteins in the cell (Figures 1K and K’).

Mammalian and *Drosophila Yap* homologues promote proliferation [21, 34]. To determine if Yap regulates proliferation in *Hydra*, we performed EdU incorporation assays on *GFP*-expressing polyps electroporated with *shGFP* alone, with *shGFP and shHyYap*, or with *shGFP and shHyYapscr*. The graph in Figure S2H shows the ratio of EdU-positive GFP^+^ cells to EdU-positive GFP^−^ cells determined for each individual polyp 5 – 6 days after electroporation. These results demonstrate that reduction of *HyYap* significantly slows the cell cycle, consistent with a role for HyYap in promoting cell proliferation.

### *HyYap* represses bud formation

Unexpectedly, knockdown of *HyYap* caused a significant increase in the number of polyps with buds (Figure 2A). All buds formed at the normal location, the budding zone. Budding normally starts with an evagination of both ectoderm and endoderm, with the epithelial cells where budding is initiated forming the tip of the bud [19]. Staining for GFP and HyYap revealed that the majority of bud tips formed from cells lacking *HyYap*, indicating that budding was initiated in cells that had lost HyYap (Figures 2B – C’).

**Figure 2.**
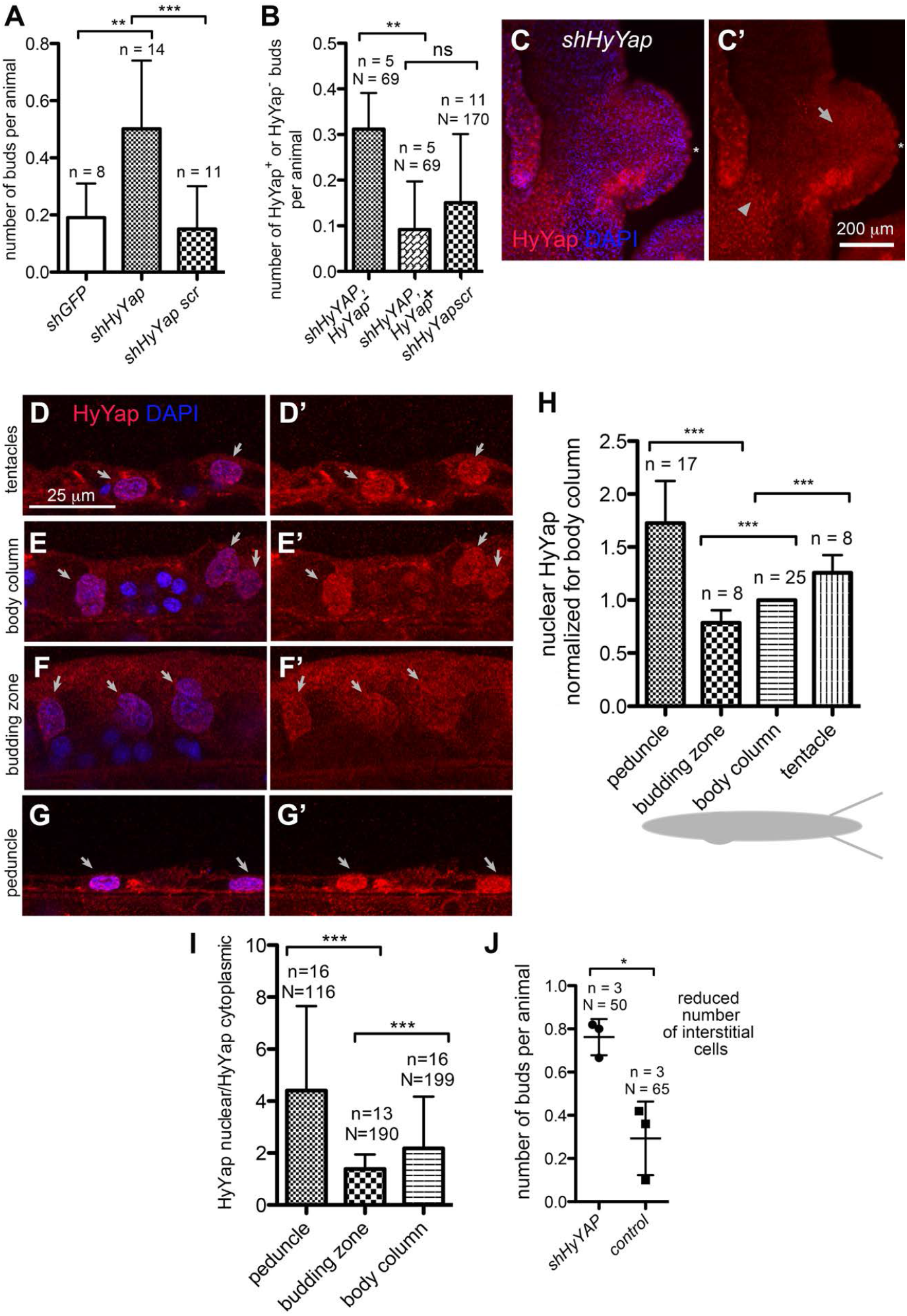
HyYap is a negative regulator of *Hydra* budding. (A) Graph showing an increased rate of budding in polyps electroported with *shHyYap* compared to controls; n – number of experiments, each experiment included 10 – 20 polyps. (B) Graph shows that a significant majority of buds developed in polyps electroporated with *shHyYap* originated from HyYap^−^ tissue; n – number of experiments, N – total number of polyps used in analysis. (C, C’) Lateral view of a bud developing from HyYap-tissue; asterisk points to the tip of the bud, arrow – to HyYap^−^ tissue, arrowhead – to HyYap^+^ tissue. (D - G’) Lateral view of the ectoderm of tentacles (D, D’), body column (E, E’), budding zone (F, F’) and a peduncle (G, G’) immunostained with anti-HyYap serum; arrows point to nuclei of ectodermal epithelial cells. (H) Graph shows the intensities of HyYap immunostaining in nuclei of tentacles, body column, budding zone and a peduncle normalized for intensity in the nuclei of a body column and superimposed on a schematic drawing of *Hydra*; for each animal, the average intensity of immunostaining was measured in the nuclei of ectodermal epithelial (10 – 20 nuclei for each area); n – number of polyps. (I) Graph shows nuclear/cytoplasmic ratio of HyYap in the body column, budding zone, and peduncle determined by immunostaining for each cell; N – number of cells, n – number of polyps. (J) Graph shows the budding rate in animals treated with 10 mM hydroxyurea for 48h prior to electroporation; n – number of experiments, N – number of polyps.

Since downregulation of *HyYap* led to bud formation, we hypothesized that budding normally occurs in areas of low HyYap expression. We quantified HyYap staining in ectodermal epithelial cells along the body column of the polyp (Figures 2D - G’). Staining intensities were measured in the nucleus and cytoplasm for each cell. Interestingly, both the lowest level of nuclear HyYap (Figure 2H) and the lowest nuclear/cytoplasmic ratio of HyYap were observed in the budding zone (Figure 2I). In contrast, the highest level of nuclear HyYap, and highest nuclear/cytoplasmic HyYap ratio were observed in tentacles and the peduncle (Figures 2H and I).

Single cell transcriptome data indicate that *HyYap* is expressed in the interstitial cell lineage [27, 35]. To determine if ectopic budding is caused by loss of *HyYap* in ectodermal epithelial cells or interstitial cells, polyps were treated with 10 mM hydroxyurea (HU) for 48h prior to electroporation. This treatment eliminates at least 50% of interstitial cells from the body column, without affecting the number of epithelial cells [36]. Increased budding was still observed upon *HyYap* knockdown (Figure 2J), implying that HyYap functions in epithelial cells to restrict budding. Together these data indicate that *HyYap* acts as a negative regulator of budding and suggested that control of the nuclear/cytoplasmic distribution of HyYap may be an important mechanism for regulating bud initiation.

### HyLATS regulates HyYAP localization and bud formation

LATS kinases phosphorylate YAP, promoting its retention in the cytoplasm and degradation (Figure 1G) [21, 22, 37]. Reduced LATS, thus, promotes active, nuclear YAP. We generated polyps mosaic for *HyLATS* knockdown by electroporating *GFP*-expressing polyps with *shHyLATS/shGFP* (Table S1). Electroporation resulted in a significant decrease of *HyLATS* mRNA (Figure S2I). Importantly, immunostaining revealed increased nuclear accumulation of HyYap in epithelial cells electroporated with *shHyLATS* and *shGFP* (Figures 3A, A’, A1, A2 and B). The nuclear/cytoplasmic ratio of HyYap was also significantly higher in cells electroporated with *shHyLATS* (Figure 3C). Thus LATS regulates Yap nuclear localization in *Hydra*, as it does in bilaterians.

**Figure 3.**
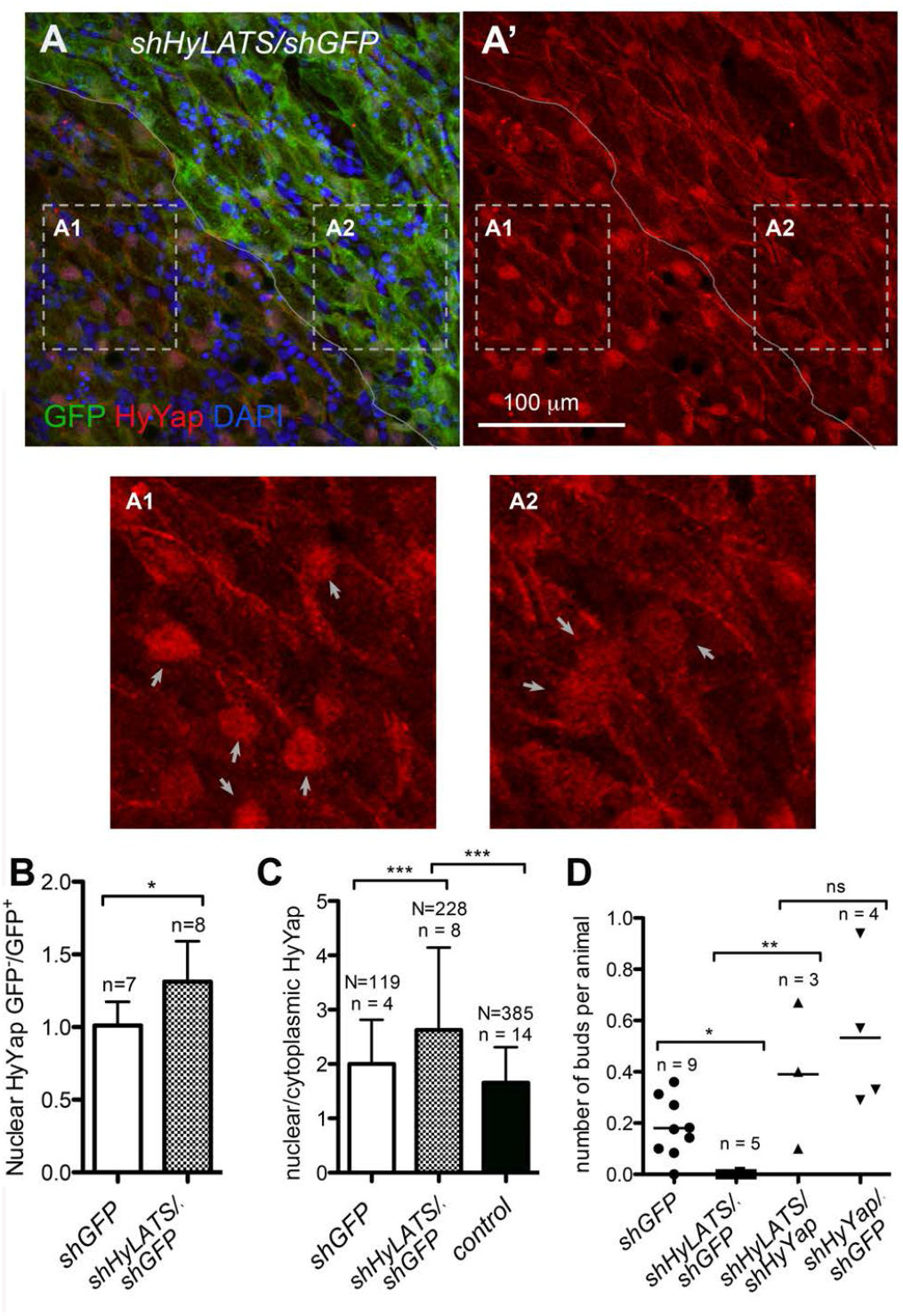
HyLATS regulates cellular localization of HyYap and the budding rate. (A, A’, A1, A2) Apical view of *Hydra* ectoderm electroporated with *shHyLATS/shGFP* and immunostained for GFP and HyYap; the border between GFP^+^ and GFP^−^ areas is marked; arrows point to nuclei; panels A1 and A2 are high magnification of GFP^−^ and GFP^+^ areas. Nuclear abundance of HyYap increases in GFP^−^ ectodermal epithelial cells. (B) Graph shows the ratio of nuclear HyYap intensities between GFP^−^ and GFP^+^ areas of *GFP* polyps electroporated with either *shGFP* or *shGFP/shHyYap*. Note, that electroporation with *shGFP* alone does affect the nuclear abundance of HyYap (GFP-/GFP+ ~ 1). The average intensities of immunostaining were measured and the ratios were calculated individually for each animal; n = number of polyps. (C) Graph shows the ratio between nuclear and cytoplasmic HyYap in non-electroporated GFP^+^ ectodermal epithelial cells (green bar), cells electroporated with *shGFP* alone, and cells electroporated with *shGFP/shHyLats*. Nuclear/cytoplasmic ratio was measured and calculated for each individual cell; N - number of cells, n – number of polyps. (D) Graphs shows the budding rate of hydras electroporated with either *shGFP* (control), or *shHyLats/shGFP*, or *shHyLats/shHyYap*, or *shHyYap/shGFP*; n – number of experiments, in each experiment 10 – 20 polyps were used in each experiment for each condition.

Since knocking down *HyLATS* resulted in increased nuclear accumulation of HyYap, we expected a concomitant increase in epithelial cell proliferation. Indeed, incorporation of EdU was higher in *HyLATS* knockdown cells than in *GFP* knockdown controls, but not significantly higher (Figure S2H). This may be explained by incomplete knockdown of *HyLATS*. Significantly, knocking down *HyLATS* halted production of buds (Figure 3D). Bud formation was rescued when animals were electroporated with a combination of *shHyYap* and *shHyLATS* (Figure 3D). These data imply that the amount of nuclear HyYap in ectodermal epithelial cells of the budding zone is a controlling factor for bud formation.

To test the effects of overexpression of nuclear HyYap, we generated a transgenic *Hydra* line that constitutively expressed a gain of function mutant *HyYap* in ectodermal epithelial cells (Figures S3A, B and C). *HyYapS72A*, bore a S72A substitution that should make it resistant to LATS phosphorylation and subsequent cytoplasmic retention and degradation [29]. The use of an operon expression construct marked *HyYapS72A*-expressing cells with DsRed2. We were able to establish a culture of mosaic (30% – 70% transgenic cells) *HyYapS72A* animals. These animals had markedly reduced budding (Figure S3D), and eventually lost the ability to propagate. Thus, loss of *HyYap* increases budding and loss of *HyLATS* or constitutive *HyYap* gain of function suppresses budding. These data indicate that the Hippo pathway acts as a regulator of bud formation.

### Inhibition of the Hippo pathway alters *Hydra* morphology

In addition to suppression of budding, knockdown of *HyLATS* had a strong effect on morphology. Tentacles containing cells in which *HyLATS* was knocked down became shorter and thicker (Figures 4A and B). We wondered if these dramatic alterations in morphology were due to overactivation of HyYap. Significantly, double electroporation of *shHyLATs* and *shHyYap* rescued the morphological defects caused by loss of HyLATS (Figure 4B) indicating that the thick tentacles were the result of excess HyYap activity.

**Figure 4.**
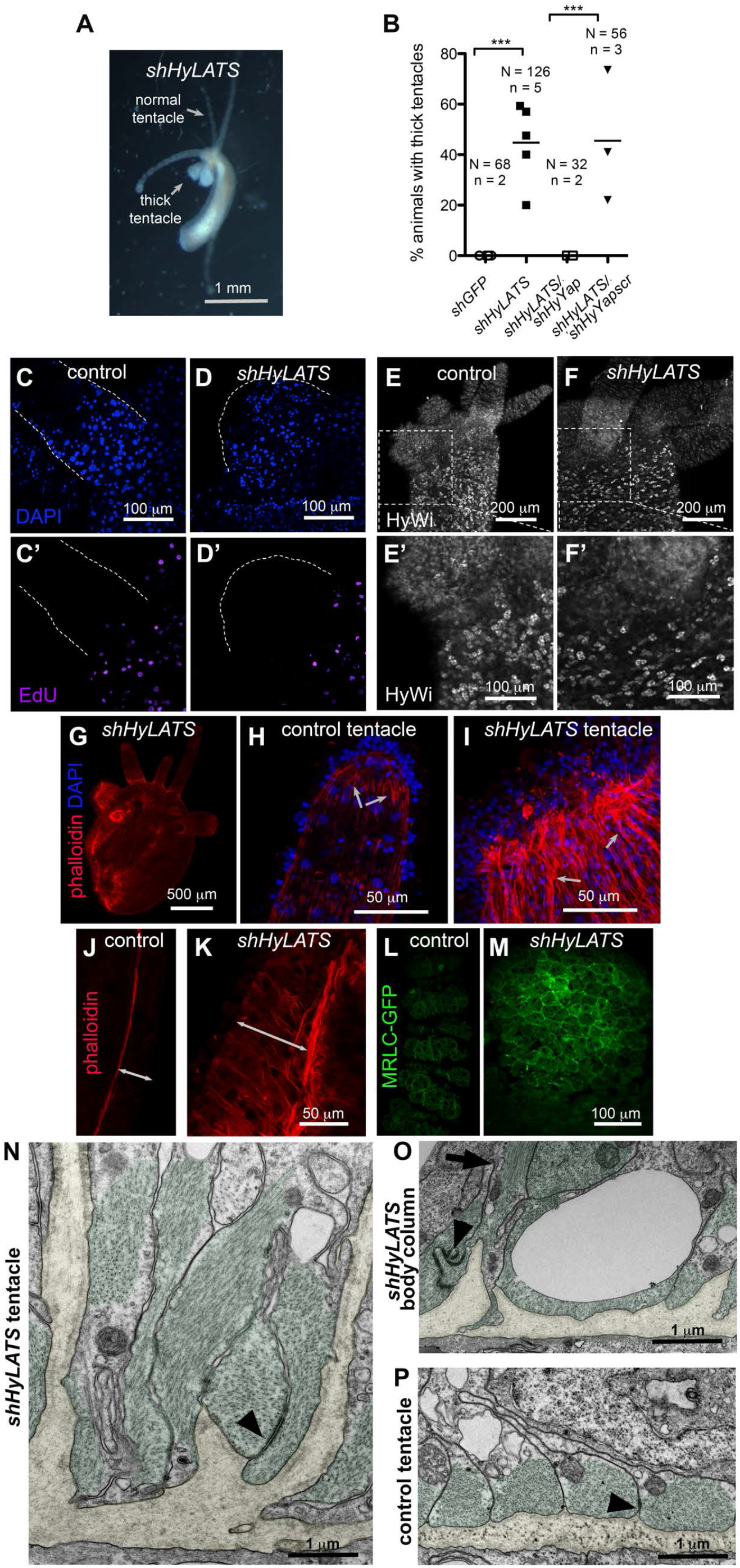
Inhibition of Hippo pathway changes *Hydra* morphology. (A) Live polyp electroporated with *shHyLats*, 14 days after electroporation. (B) Graph shows the percentage of thick tentacle formation in polyps electroporated with *shGFP* (control), *shHyLATS* alone, *shHyLATS/shHyYap* and *shHyLATS/shHyYapscr*; N – number of polyps, n – number of experiments. (C – D’) EdU is not detected in either control (C, C’) or thick (D, D’) tentacles. Edges of tentacles are outlined by dotted lines. (E – F’) The stem/progenitor marker HyWi is not detected in either control (E, E’) or thick (F, F’) tentacles. (G) *Hydra* polyp electroporated with *shHyLATS* and stained with phalloidin, 12 days after electroporation. Visible tear along the body column is an artifact of fixation and is common when shortened *shHyLats* – electroporated polyps are fixed. (H, I) Ends of normal (H) and thick (I) tentacles stained with phalloidin. Arrows point to the ectodermal muscle processes that are filled with actin fibers and oriented along the length of the tentacle. (J, K) Lateral view of the ectoderm of the body columns underneath normal tentacle (J) and thick (K) tentacle stained with phalloidin. Double-headed arrows indicated the thickness of the ectoderm (L, M) Apical view of the normal (L) and thick (M) tentacles of *MRLC-GFP* polyps immunostained for GFP. (N - P) Transmission electron microscopy of cross sections visualizing the basal compartment of ectodermal epithelial cells in *shHyLats* tentacles (N), shHyLats body column (O), and wild type tentacles (P). (N, O) Muscle processes (green) exhibit abnormal elongation along the apical-basal axis of the cells and sometimes ectopic positioning distant from the mesoglea. Muscle processes remain connected by normal numbers of spot desmosome-like junctions (arrowheads), but they show a dramatic loss of their parallel alignment along the polyp’s oral-aboral body axis as shown in wild type controls (P). (N, O) At positions, where the disrupted planar array of muscle fibers had gaps, the mesoglea (yellow) folded into the cytoplasm of the ectodermal epithelial cells without losing the hemidesmosome-like junctions usually located at the basal membrane surface of epithelial cells. (O) shows a representative image with an ectopic muscle process running along the apical-basal axis (arrow) pointing toward the mesoglea folding.

The epithelial cells of the body column of *Hydra* are continually displaced into the tentacles, where they arrest in the G2 phase of the cell cycle and terminally differentiate (Figure 1B) [10]. EdU labeling indicated LATS knockdown cells still underwent G2 arrest. *HyWi*, the *Hydra* homologue of *piwi* [11, 38] marks undifferentiated epithelial cells (i.e. cells that have not been displaced into the tentacles or the basal disk). Immunostaining with HyWi antibodies showed no change compared to controls (Figures 4C - F’), indicating LATS knockdown did not prevent formation of differentiated cells.

Examination of the thick tentacles and the body column below them revealed thickening of the ectoderm (Figures 4G, H and I), an accumulation of ectopic actin fibers (Figures 4 H - K), and accumulation of myosin along the apical surface of ectodermal epithelial cells (Figures 4L and M).

Consistent with the results of *HyLATS* knockdown, mosaic *HyYa*p*S72A* transgenic polyps also often developed short thick tentacles (Figure S3B). *HyYapS72A* transgenic cells were elongated in the apico-basal direction and had ectopic actin fibers (Figures S3C, E, F and G) as in *HyLATS* knockdown polyps. These data support the proposal that HyLATS and HyYAP act together in a pathway to regulate body morphology. Eventually, cells expressing *HyYapS72A* took over the animal and due to morphological defects the animals were unable to feed themselves and died.

To understand the cellular basis for the altered morphology, we analyzed ultra-thin sections of *shHyLATS* thick tentacles and adjacent body column tissue using transmission electron microscopy. While the apical compartment of ectodermal epithelial cells and the endodermal layer did not exhibit obvious defects in cytoplasmic organization, there was a dramatic loss of normal structure in the basal part of ectodermal epithelial cells (Figures 4 N - P, Figure S4). *HyLATS* knockdown affected shape, positioning, and orientation of muscle processes. We observed gaps in the planar array of the processes, individual processes elongated along the cells apical-basal axis, and some processes detached from the mesoglea.

The parallel alignment of neighboring processes was strongly disrupted and randomized, sometimes even the parallel alignment of actin filaments within a muscle process was lost (Figures 4N and O). There was a clear loss of planar polarity in this tissue layer. Interestingly, the mesoglea adjacent to the *shHyLATS* knockdown cells showed folding towards the apical surface of the ectoderm (Figures 4 N and O, Figures S4C and D). In the body column, we detected muscle processes apical to the tip of mesoglea folding running perpendicular to their normal planar orientation (Figure 4O). Thus, disruption of the normal, non-folded mesoglea sheet may be a result of contraction of these ectopic muscle fibers along the epithelial cell’s apical-basal axis. *shHyLATS* tentacle and body column tissue both showed these defects, but knockdown phenotypes were clearly stronger in the tentacles than in the body column.

### The Hippo pathway acts upstream of the canonical Wnt pathway during budding

The canonical Wnt pathway is activated early in budding, and induces axis formation in *Hydra* [4, 6, 14]. Experimental activation of the Wnt pathway in the *Hydra* body column leads to induction of ectopic axes, and inhibition of Wnt signaling blocks axis formation [4, 16]. Since knocking down of *HyYap* led to increased bud formation, we hypothesized that the Hippo pathway lies upstream of Wnt/β-catenin signaling. In early buds, *HyWnt3* expression is seen in a patch of 15-20 cells, soon after thickening of the ectoderm and expression, and remains at the tip of the growing bud [14] (Figures 1E and 5A). To test the effects of the Hippo pathway on canonical Wnt signaling, we first examined expression of *HyWnt3* upon loss of *HyYap*. Significantly, more polyps electroporated with *shHyYap* had *HyWnt3* patches in the budding zone than controls (Figure 5B).

**Figure 5.**
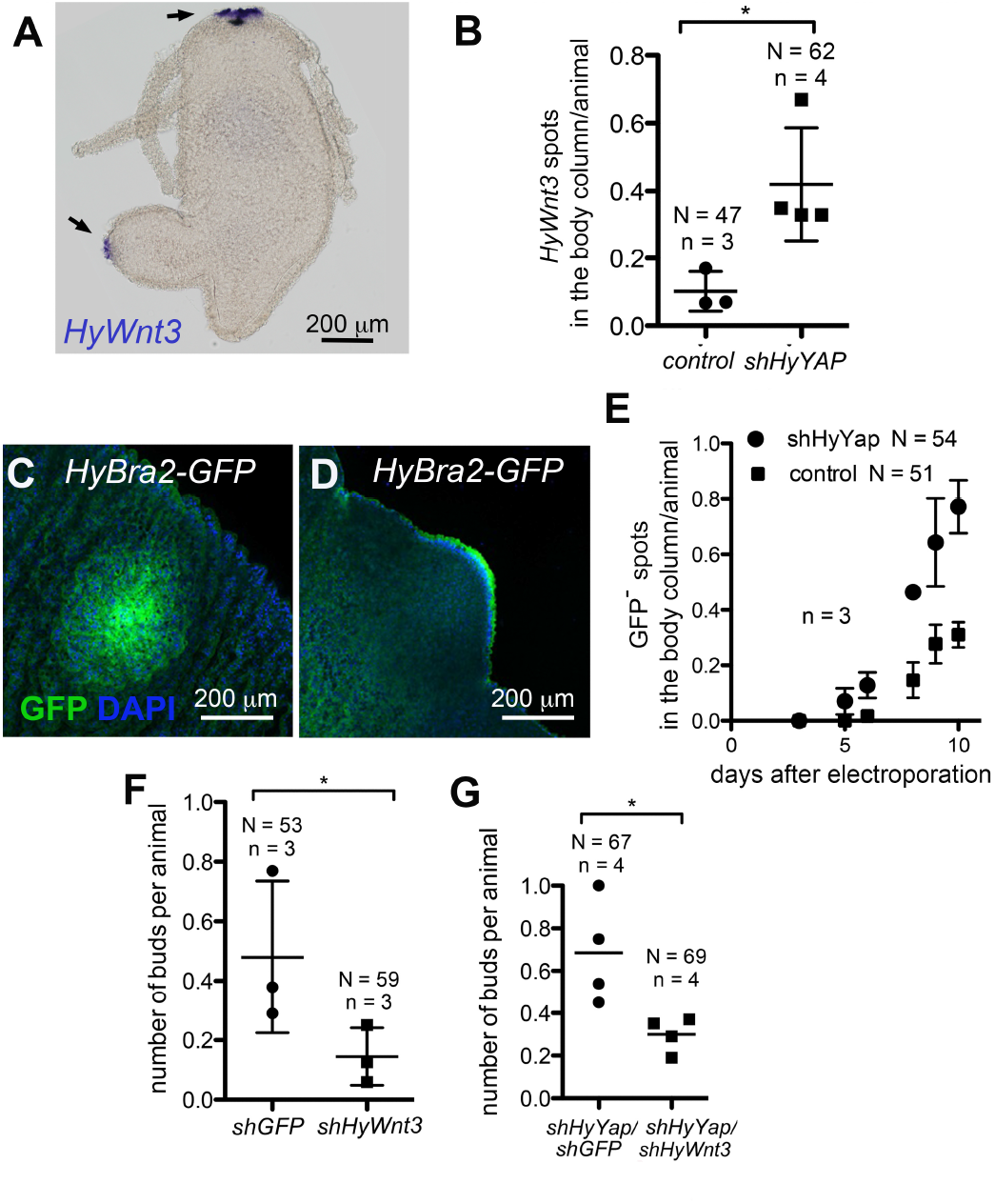
*HyYap* acts upstream of canonical Wnt signaling during budding. (A). Whole mount *in situ* hybridization of *Hydra* with anti-*HyWnt3* probe. Arrows point to areas of *HyWn3t* expression. (B) Graphs shows the increased number of *HyWnt3* spots in the body column of polyps electroporated with *shHyYap*. (C,D) Anti-GFP immunostaining of *HyBra2-GFP* polyps electroporated with *shHyYap*, 6 days after electroporation. (C) – apical view, (D) – lateral view of GFP^+^ bud. (E) Graph shows the increased number of GFP^+^ spot in the body column of *HyBra2-GFP* polyps electroporated with *shHyYap*. (F) Graph shows decreased budding rate in polyps electroporated with *shHyWnt* compared to hydras electroporated with *shGFP*. N – number of animals, n – number of experiments. (G) Graph shows the decreased budding rate in polyps electroporated with *shHyYap/shHyWnt* compared with hydras electroporated with *shHyYap* alone. We also examined expression of *HyBra2*, a hypostome-specific gene, which is induced by Wnt expression [39]. *HyBra2* is expressed early in budding and expression persists in the hypostome of the growing bud [39]. We used a transgenic *Hydra* line that expresses *GFP* under the control of the *HyBra2* promoter [40] to assay induction of budding upon loss of HyYap (Figures 5C and D). GFP patches were observed in the budding zone in a significantly higher number of animals upon electroporation with *shHyYap* compared to controls (Figure 5E), indicating that *HyBra2* expression is activated upon loss of *HyYap*.

Knockdown of *shHyWnt3* alone results in reduced production of buds, as expected from the key role of *HyWnt3* in axis initiation [41](Figure 5F). To test if budding by loss of *HyYap* is mediated by increased *HyWnt3*, we simultaneously knocked down *HyYap* and *HyWnt3*. Importantly, the increased budding that resulted from downregulation of *HyYap* was suppressed when *shHyWnt3* was electroporated along with *shHyYap* (Figure 5G). These data indicate that HyYAP negatively regulates HyWnt3 expression to suppress bud formation in *Hydra*. The Hippo pathway thus integrates growth and axis formation by controlling Wnt signaling.

## Discussion

The canonical Wnt (Wnt/β-catenin) pathway is both necessary and sufficient for axis formation in cnidarians: transplantation of the hypostome, the organizer, into a body column of a host, or experimental induction of canonical Wnt signaling leads to formation of a new axis [4, 5, 7]. However, in *Hydra*, the onset of *HyWnt3* expression occurs after the first signs of bud formation [14] pointing to events regulating budding upstream of *HyWnt3*. Early studies connected budding to the cell cycle and cell displacement in *Hydra* [10, 11, 42], however, molecular mechanisms connecting these processes were never illuminated.

In a normally fed *Hydra* polyp, the rate of cell division is similar along the body column [11]. Due to continuous cell division, cells below the subtentacle zone are being pushed down the body column [11]. In the budding zone, excess cells are forced out of the maternal axis to form a new axis, a bud [10]. Buds do not form in starved *Hydra*, since the rate of cell division can not compensate for the rate of cell loss at the ends [13]. Ectodermal epithelial cells that are about to form a bud are packed tighter than cells of the body column: they elongate in apico-basal direction and reduce the area of attachment to mesoglea, the basement membrane (Fig. 1F) [20]. Nuclear localization of mammalian and *Drosophila* Yap is negatively regulated by cell density [22, 29]. Similar, we find that in *Hydra*, highly packed cells of the budding zone have less nuclear HyYap than body column cells, and, especially than the flat cells of the peduncle and tentacles (Figures 2 and 6A). Since downregulation of *HyYap* leads to induction of budding, we suggest that HyYap might act as a molecular link between continuous cell division and budding (Figure 6B). A seeming paradox is that loss of HyYap reduces proliferation. However, knocking down of *HyYap* does not completely block epithelial cell division. In addition, when *HyYap* is knocked down locally, the rest of the body column cells continue to divide, pushing electroporated cells into the budding zone and aboral end.

**Figure 6.**
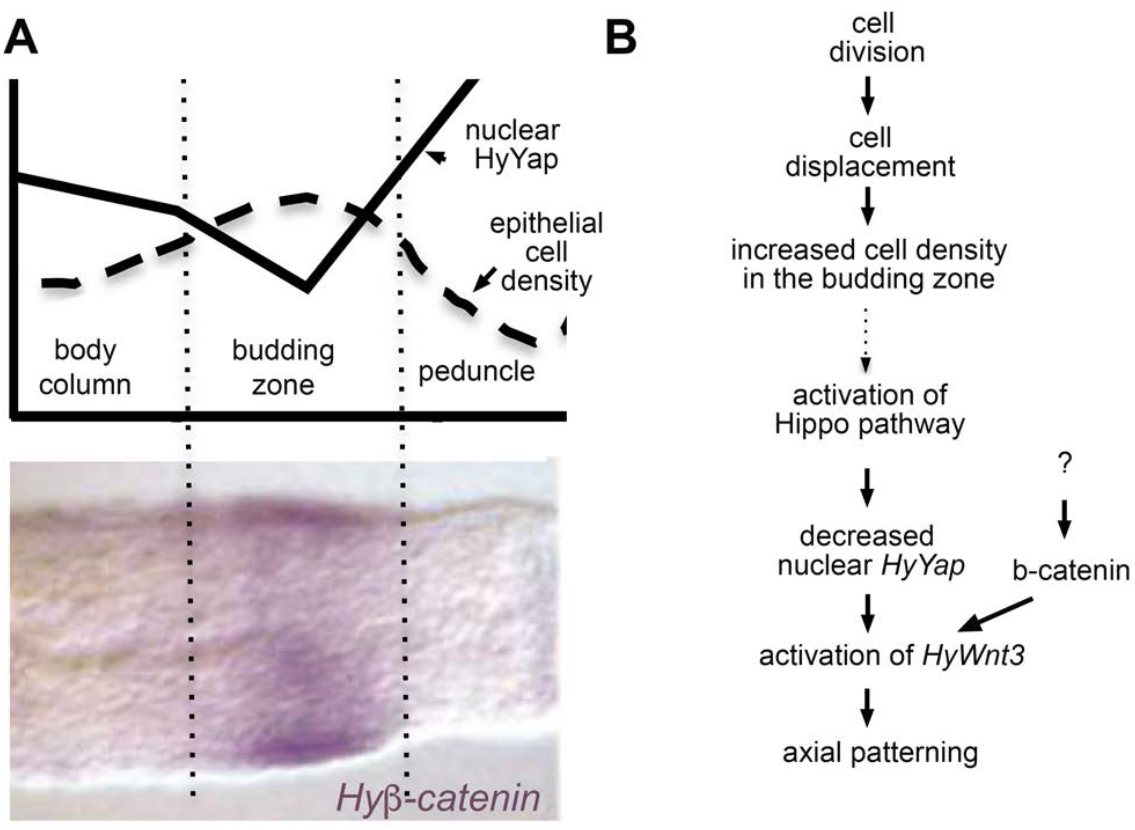
The proposed role of the Hippo pathway in axis formation in *Hydra*. (A) Schematic drawings of the axial distributions of epithelial cell density (adapted from [11]) and nuclear HyYap superimposed over *Hyβ-catenin in situ* (adapted from [14]). (B) Hypothetical model identifying HyYap as a molecular link between cell division and axis formation in *Hydra*. Dotted arrow indicates causal connection based on analogy with *Drosophila* and mammals that yet has to be demonstrated in *Hydra*.

We show that the Hippo pathway regulates budding by attenuating canonical Wnt signaling. Yap and its homologues Yorkie and TAZ also bind and inhibit Wnt signaling by binding Disheveled [43, 44]. Yap and TAZ also bind and inhibit β-catenin [45]. Interestingly, buds that were induced by knocking down of *HyYap* always formed at the normal location, the budding zone, in spite of the larger knockdown area. Intriguingly, expression of *Hyβ-catenin* is also higher in the budding zone than in surrounding body column tissue (Figure 6A) [14]. *Hyβ-catenin* can directly activate transcription of *HyWnt3* [15]. Thus, induction of *HyWnt3* through release of *Hyβ-catenin* upon knocking down of *HyYap* is a possible scenario. Also required for budding is noncanonical Wnt signaling. Noncanonical Wnt activation occurs in the densely packed ectodermal cells early in budding, before the onset of *HyWnt3* expression, and depends on *Hyβ-catenin* [16]. Whether noncanonical Wnt signaling operates downstream of Hippo or in a parallel pathway is an intriguing subject for future research.

Inhibition of the Hippo pathway by knocking down *HyLATS* results in dramatic shortening and thickening of tentacles. IF and EM analyses show that *HyLATS* knockdown leads to major changes in the actin cytoskeleton and epithelial cell shape. Interestingly, inhibition of Wart (LATS) in *Drosophila* stimulates polymerization of F-actin in a Yorkie-independent manner, by acting on the capping protein [46, 47]. In contrast, in mammalian cells, activation of F-actin requires the transcription activity of Yap [48]. In *Hydra*, knocking down of *HyYap* rescues the *shLATS* phenotype, suggesting a role for HyYap transcriptional control of actin. Abnormal polymerization of actin could explain the randomized orientation of the muscle processes and misfolding of mesoglea seen in the *HyLATS* knockdown.

To summarize, we show that the conserved LATS-Yap-Hippo signaling pathway plays a major role in morphogenesis in the cnidarian *Hydra*, and acts upstream of canonical Wnt signaling and expression of *Wnt3* during axis initiation. Our findings demonstrate that the amount of nuclear Yap decreases in the budding zone due to Hippo activation induced by high cell density in this zone, and that reduction of nuclear Yap leads to activation of Wnt/β-catenin signaling and budding (Figure 6B). We speculate that Hippo signaling, and nuclear Yap serve to link continuous cell division, cell density, and axis formation early in metazoan evolution (Figure 6B).

## Acknowledgements

We thank Joe Culotti (LTRI, Toronto) for invaluable support of the project and help in writing the manuscript, Leonid Brown (University of Guelph) for help with culturing *Hydra*, Marina Gertsenstein (Toronto Center for Phenogenomics) for help with the electroporation procedure, Thomas Bosch (University of Kiel) for providing *GFP* transgenic *Hydra*, Taylor Skokan (Universoty of California, San Francisco) for providing the *MRLC-GFP* transgenic *Hydra*, Celina Juliano (University of California, Davis) for providing anti-HyWi antibody.

## Supplemental Information

**Table S1.**
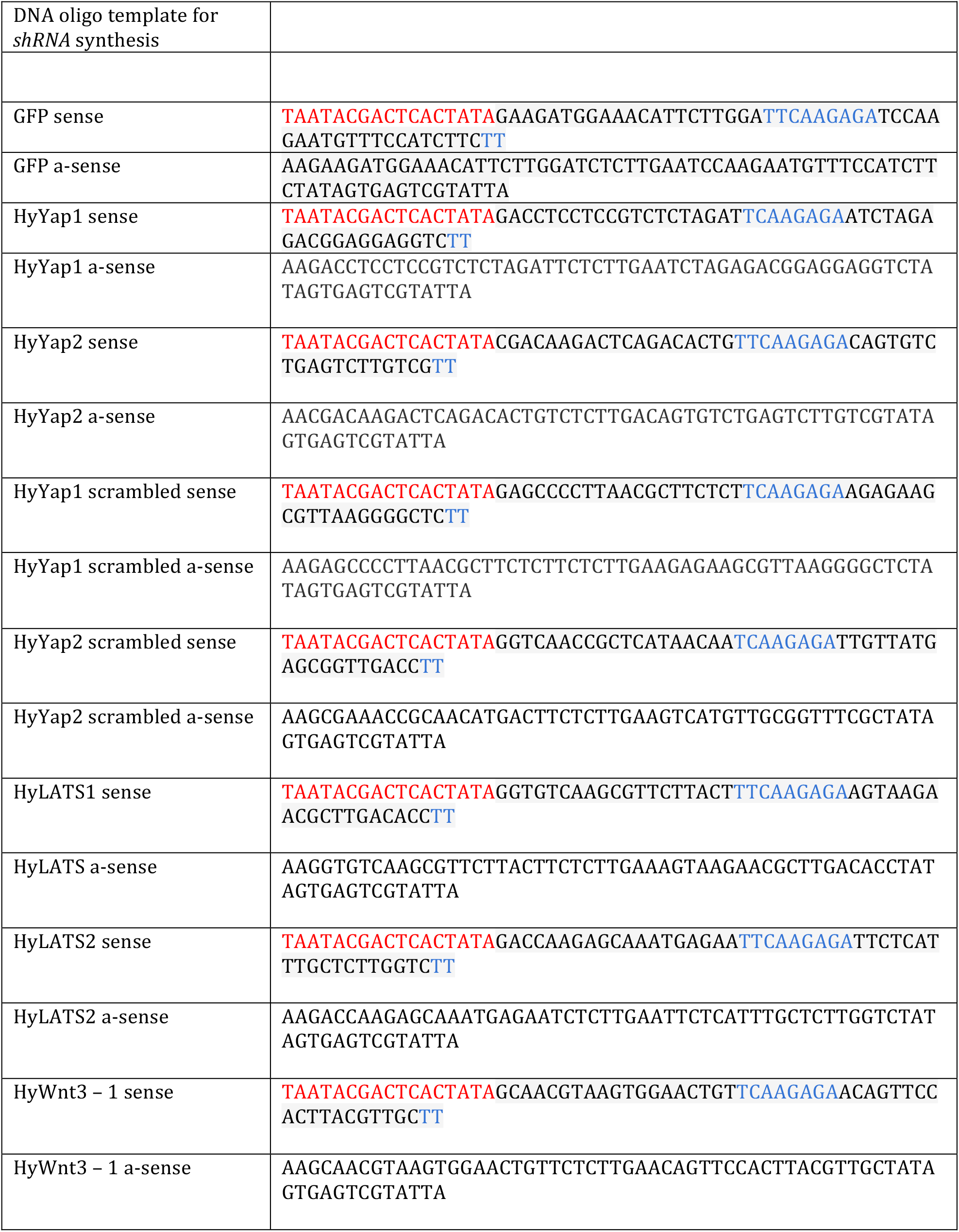
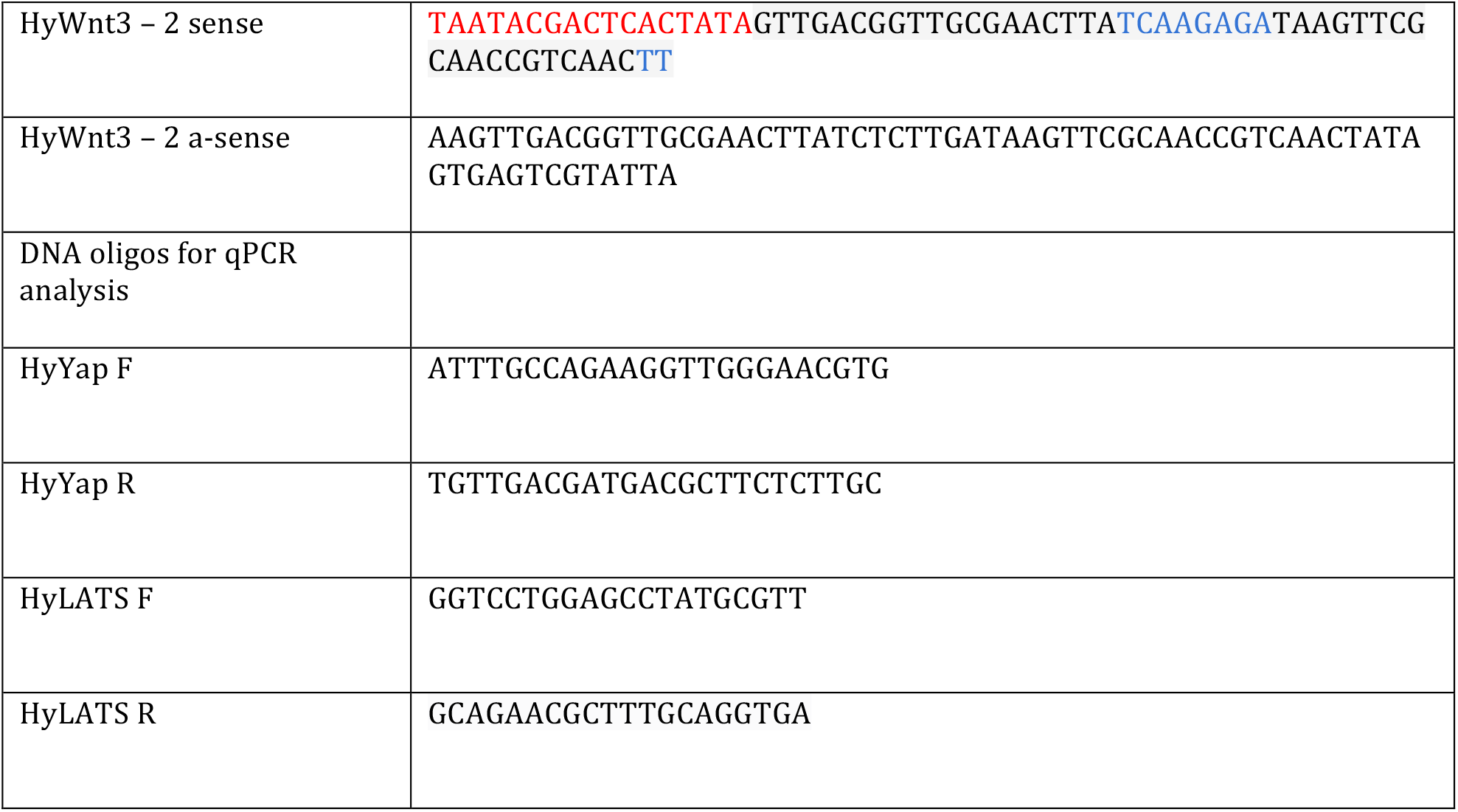
DNA oligonucleotides used in the study. T7 promoter sequence is in red, gap sequence is in blue.

**Figure S1.**
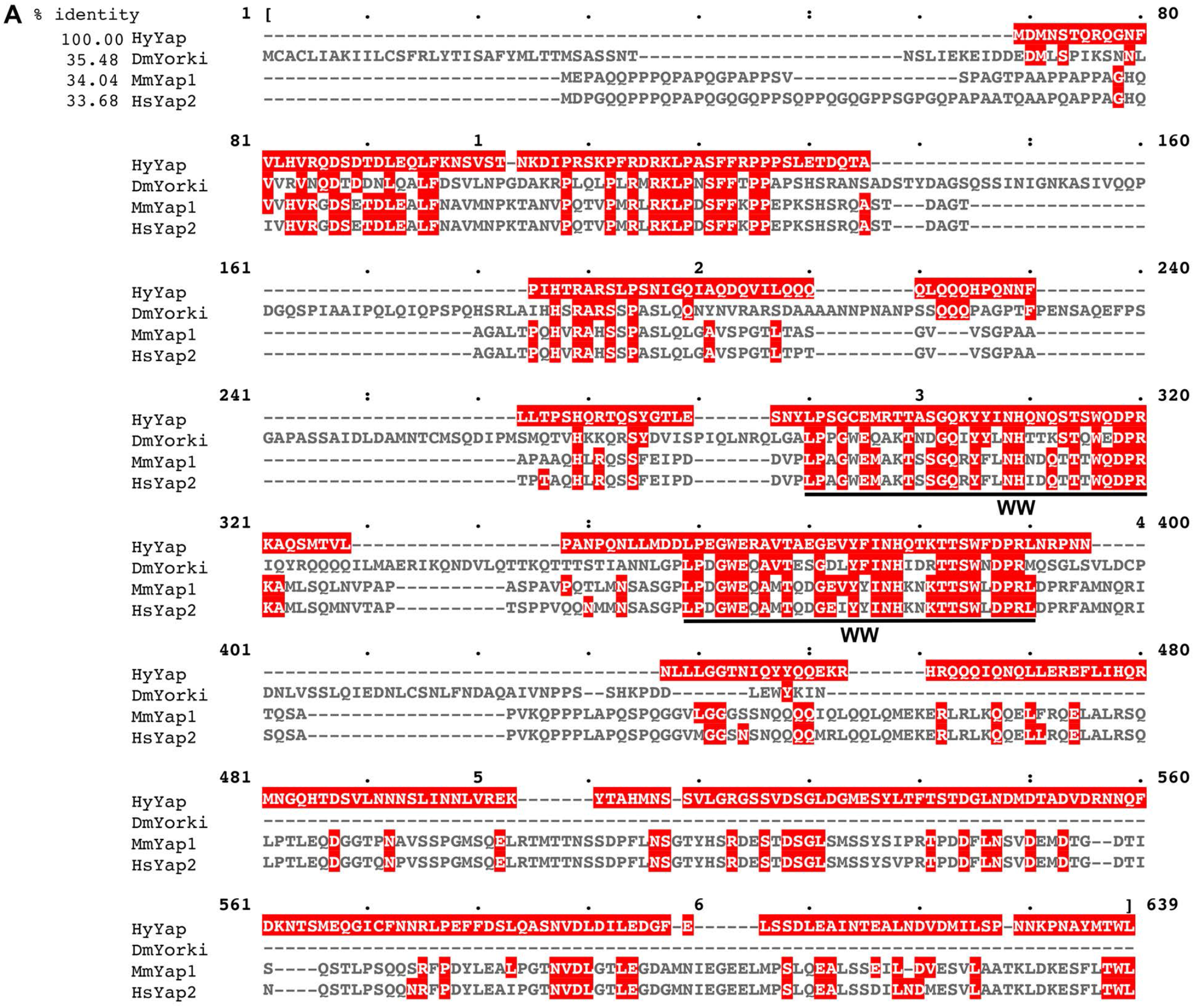
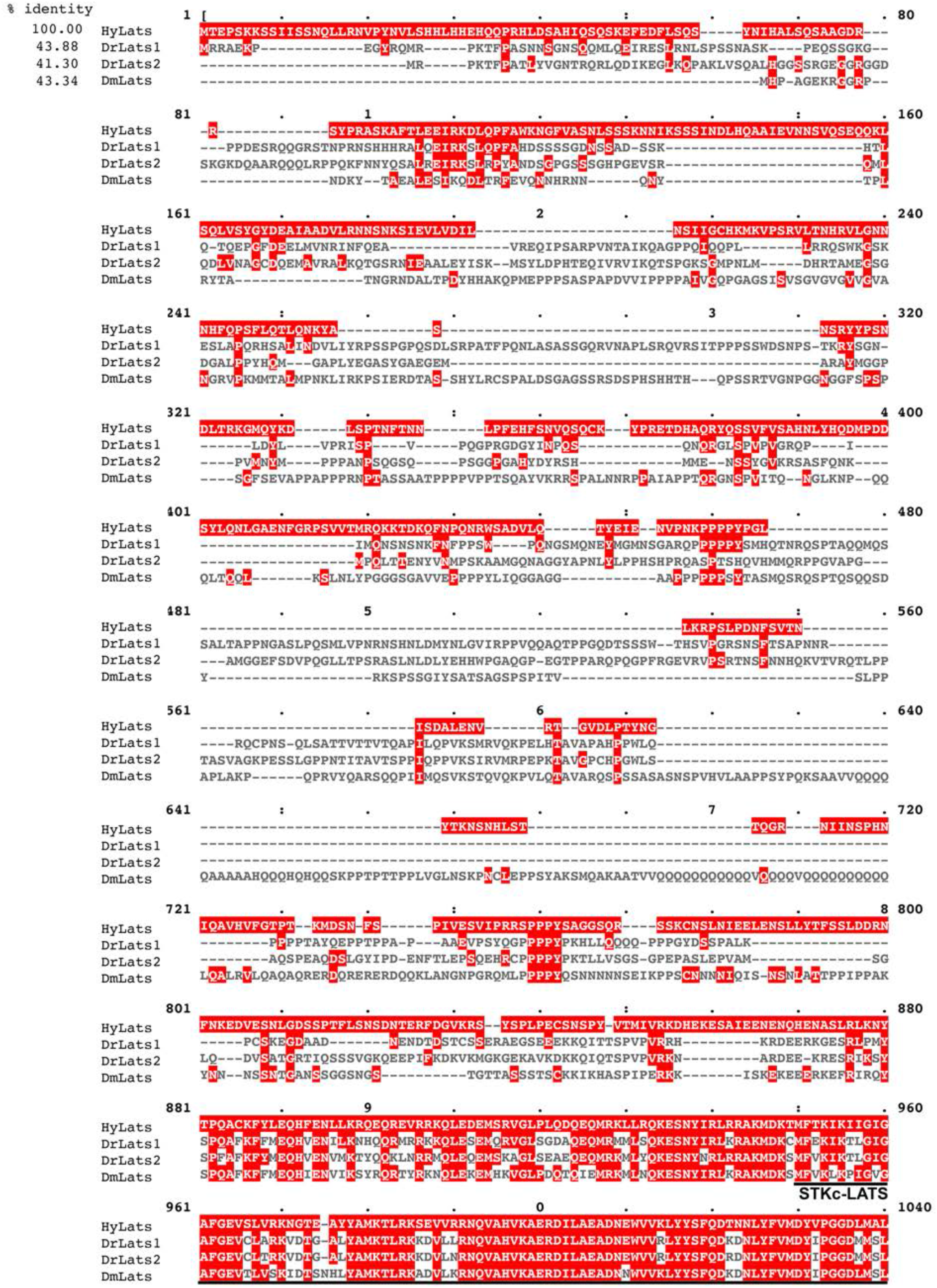
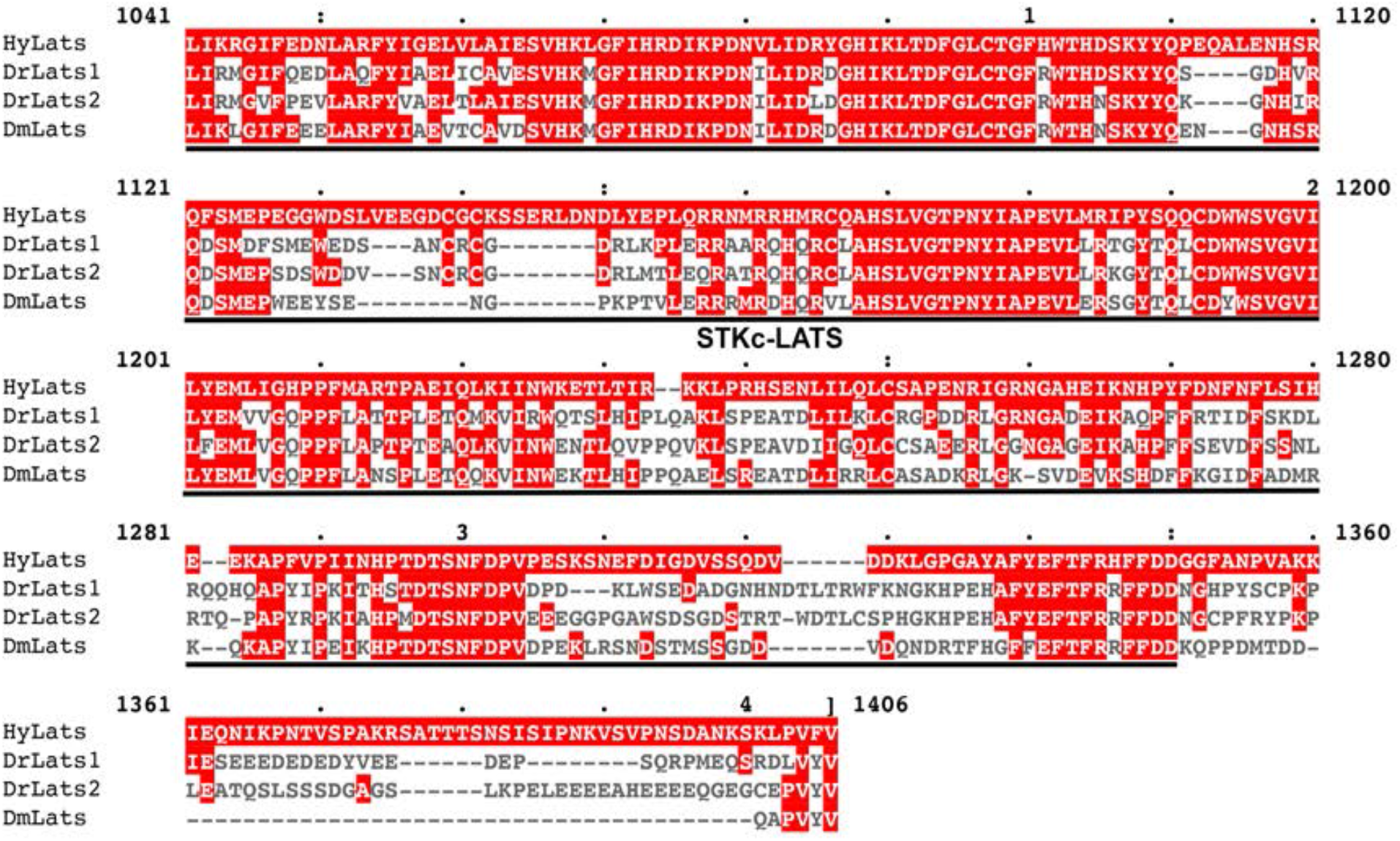
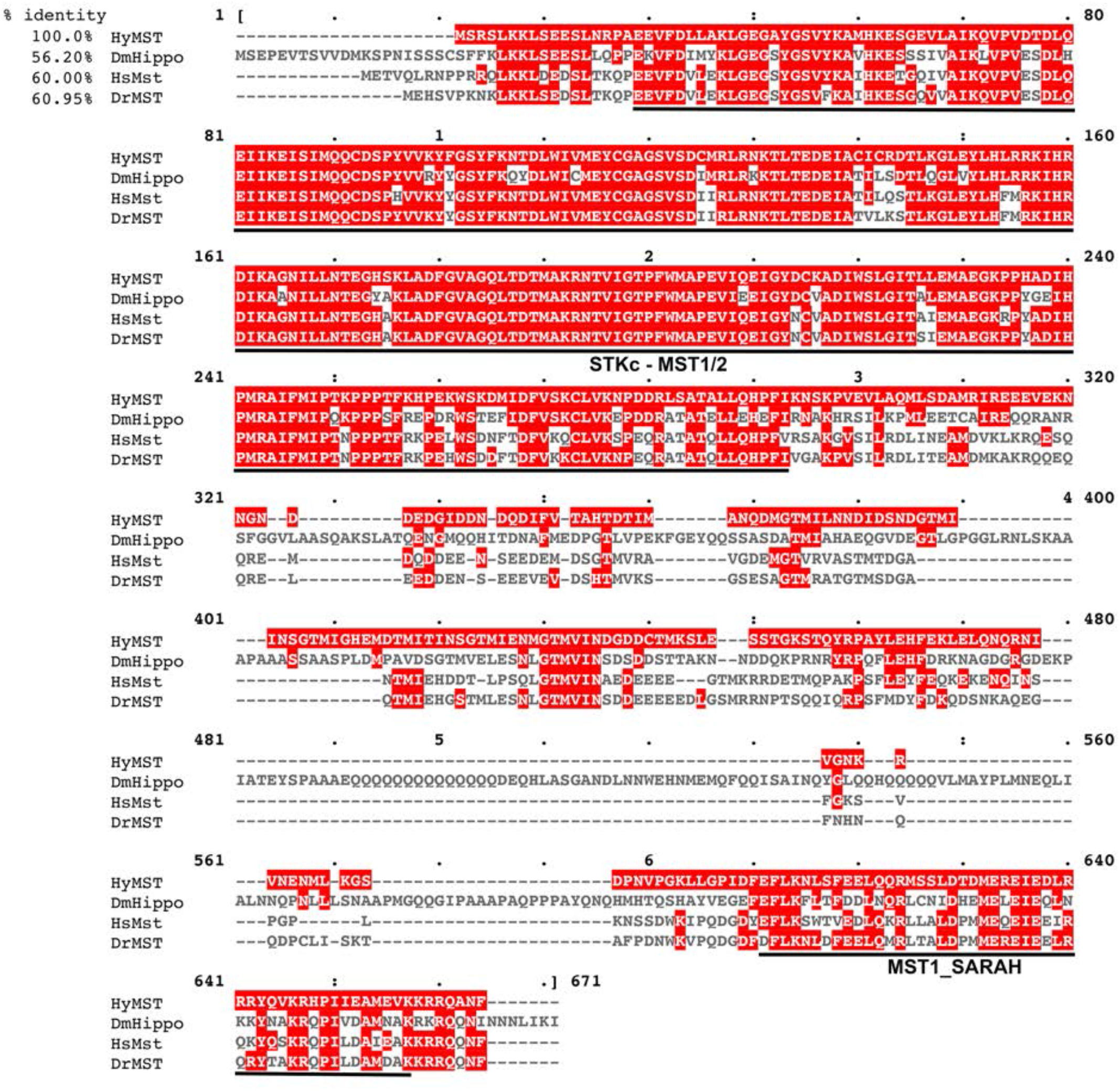
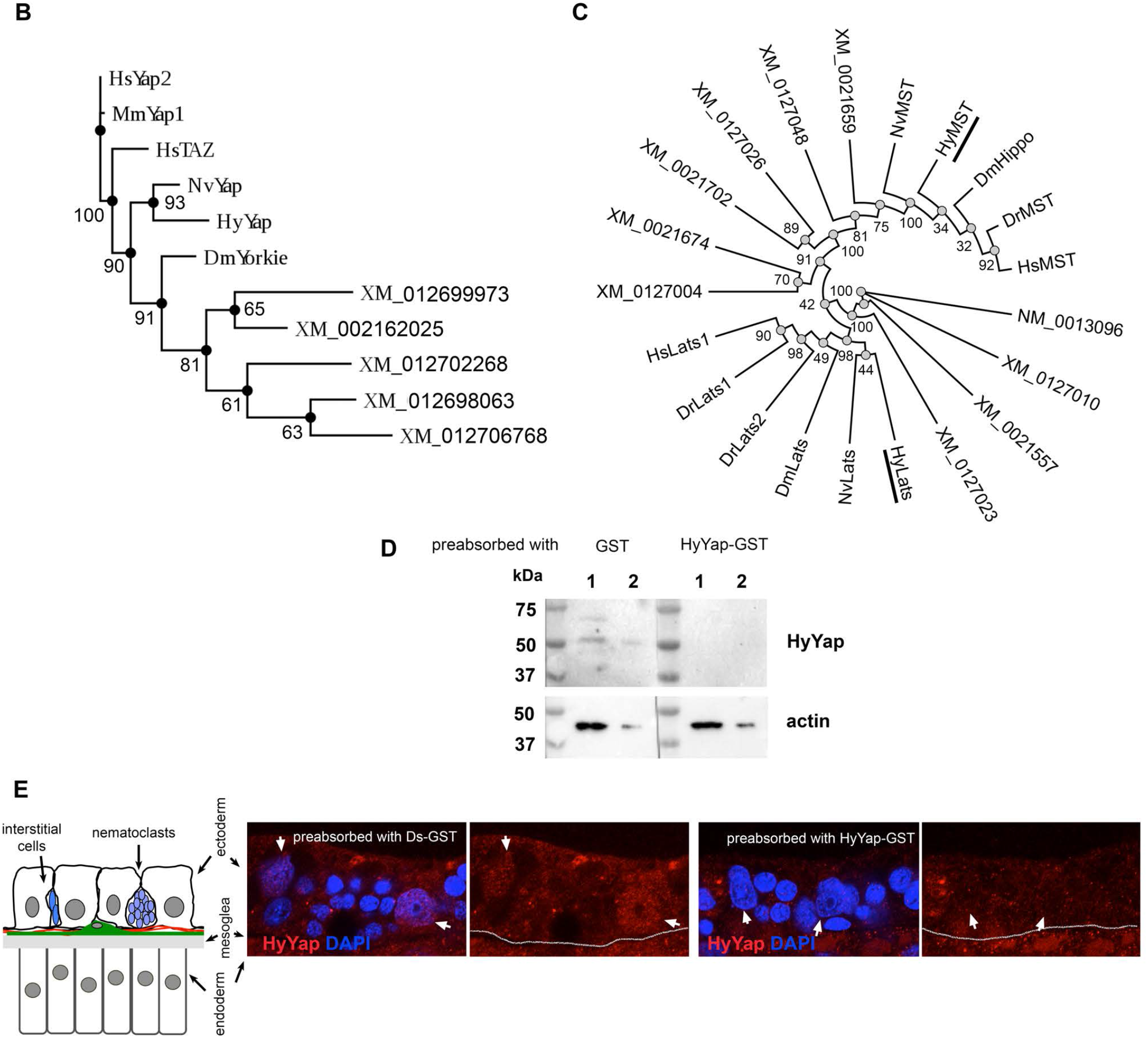
(A) Alignments of predicted *Hydra* Yap, LATS and MST proteins with *Drosophila* and vertebrate homologues. Characteristic domains of proteins are underlined: WW – tryptophan containing domain that binds specific proline-rich sequences, STKc–LATS – catalytic domain of LATS family of serine/threonine kinases, STKc–MST1/2 catalytic domain of MST family of serine/threonine kinases, MST1-SARAH – apoptosis-mediating SARAH domain of MST1 proteins. Dm – *Drosophila melanogaster*, Mm – *Mus musculus*, Hs – *Homo sapience*, Dr – *Danio rerio*. (B) Phylogenetic tree of Yap family proteins based on the complete protein sequence, Maximum Likelihood analysis (100 bootstrap replicates, bootstrap values are indicated for each node); predicted *Hydra* homologue of Yap is underlined; Hs – *Homo sapiens*, Mm – *Mus musculus*, Nv – *Nematostella vectensis*, Hy – *Hydra vulgaris*, Dm – *Drosophila melanogaster*; XM_0126999, XM_0021620, XM_0127022, XM_0126980, XM_0127067 - sequences that came at the top of BLAST search of *Hydra* nucleotide databases for Yap; sequences were aligned with Clustal Omega (https://www.ebi.ac.uk/Tools/msa/clustalo/) and analyzed using Akaike Information Criterion (http://www.atgc-montpellier.fr). (C) Phylogenetic tree of LATS and MST families based on the complete protein sequence, Maximum Likelihood analysis (100 bootstrap replicates, bootstrap values are indicated for each node); predicted *Hydra* homologues LATS and MST proteins are underlined; XM_012702323, NM_001309671, XM_012701027, XM_002155754, XM_002167407, XM_012700498, XM_002165989, XM_012704879, XM_002170224, XM_012702641, sequences that came at the top of BLAST search of *Hydra* nucleotide databases for Yap, LATS and MST are included along with predicted homologues; sequences were aligned with Clustal Omega (https://www.ebi.ac.uk/Tools/msa/clustalo/) and analyzed using Akaike Information Criterion (http://www.atgc-montpellier.fr). (D) Western blot analysis of total *Hydra* lysates with anti-HyYap serum preabsorbed with either GST or HyYap-GST. Lysates are made from either 3 animals (lanes 1) or 1 animal (lanes 2). For the loading control the same blot was immunostained with actin antibodies. (E) Lateral view of *Hydra* ectoderm immunostained with anti-HyYap serum preabsorbed with HyDs-GST or with HyYap-GST antigens. Arrows point to the nuclei of ectodermal epithelial cells; schematic drawing of the lateral view of *Hydra* ecto- and endoderm is shown on the left.

**Figure S2.**
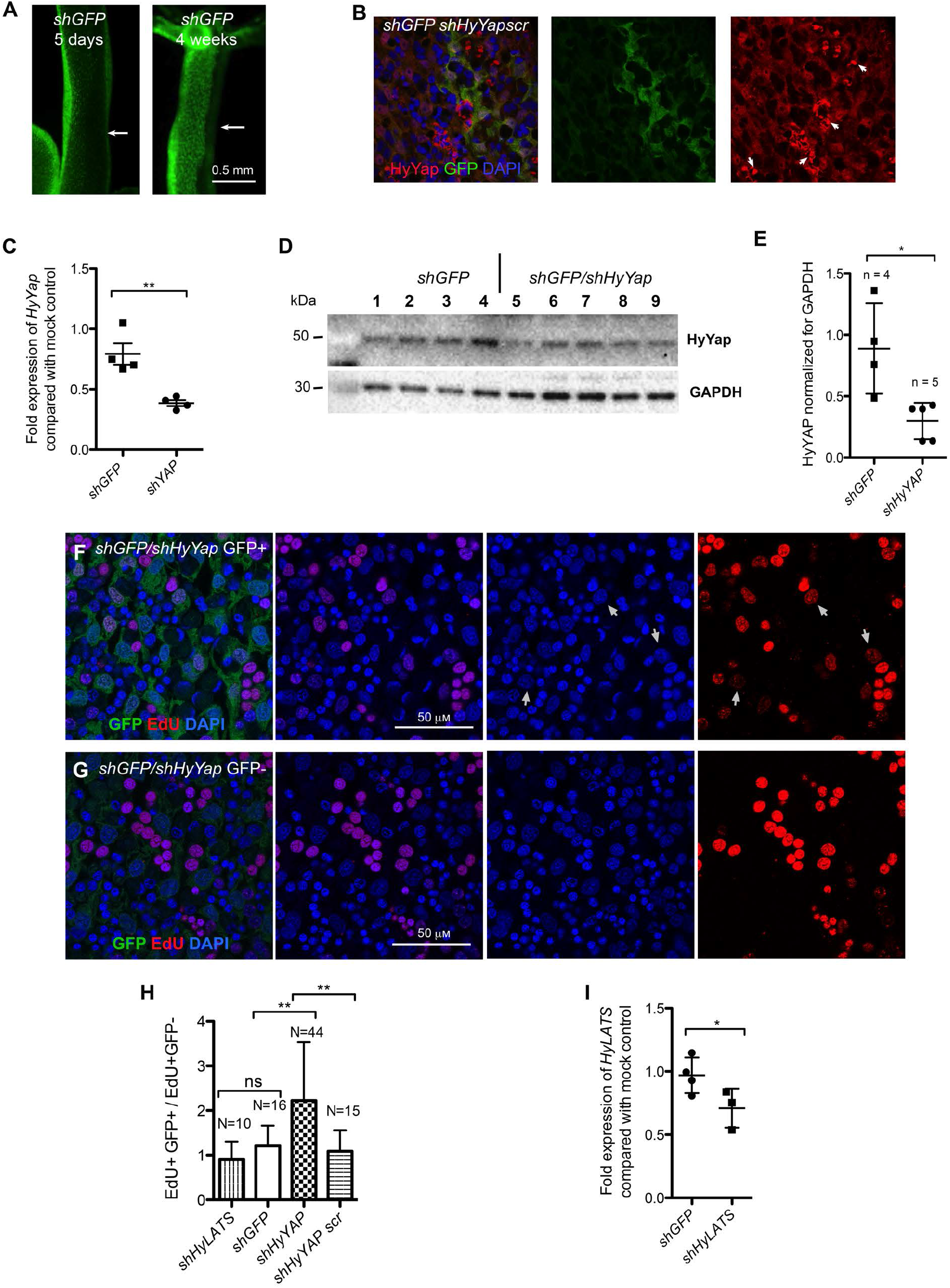
(A) *GFP* hydras electroporated with *shGFP* 5 days and 4 weeks after electroporation. Arrows point to the areas of GFP knockdown. (B) Apical view of ectoderm of *GFP* hydras electroporated with *shGFP/shHyYapscr* hairpin and immunostained with anti-GFP and anti-HyYap antibodies. Arrows point to a non-specific staining of nematocyte capsules that is observed for a variety of antibodies when used *Hydra* immunostaining protocol (C) qPCR analysis of *HyYap* in *GFP* hydras electroporated with either *shGFP* or *shGFP/shHyYap*. 4 animals for each condition. (D,E) Western blot analysis of total lysate from *GFP* hydras elecroporated with either *shGFP* alone or *shGFP/shHyYap*, 1 animal per lane; n – number of animals. For the loading control the same blot was immunostained with GAPDH antibodies. (F,G) Apical view of ectoderm of *Hydra* ectoderm with *shGFP/shHyYap*, pulse labeled with EdU 6 days after electroporation and immunostained for GFP and EdU; (F) – GFP^+^ area, (G) – GFP^−^ area; arrows point to EdU+ epithelial cells. (H) Graph shows ratio between numbers of EdU+ ectodermal epithelial cells in GFP^+^ and GFP^−^ areas in *GFP* hydras electroporated with either *shGFP* alone, *shGFP/shHyYap* or *shGFP/shHyYap scr* and pulse-labeled with EdU 6 – 7 days after electroporation; EdU+GFP+/EdU+GFP-ratio was calculated for each individual animal; the total of 60 – 200 cell was used in analysis of each animal; n – number of animals. (I) qPCR analysis of *HyLATS* in *GFP* hydras electroporated with either *shGFP* or *shGFP/shHyLATS*. Each point represents an individual animal.

**Figure S3.**
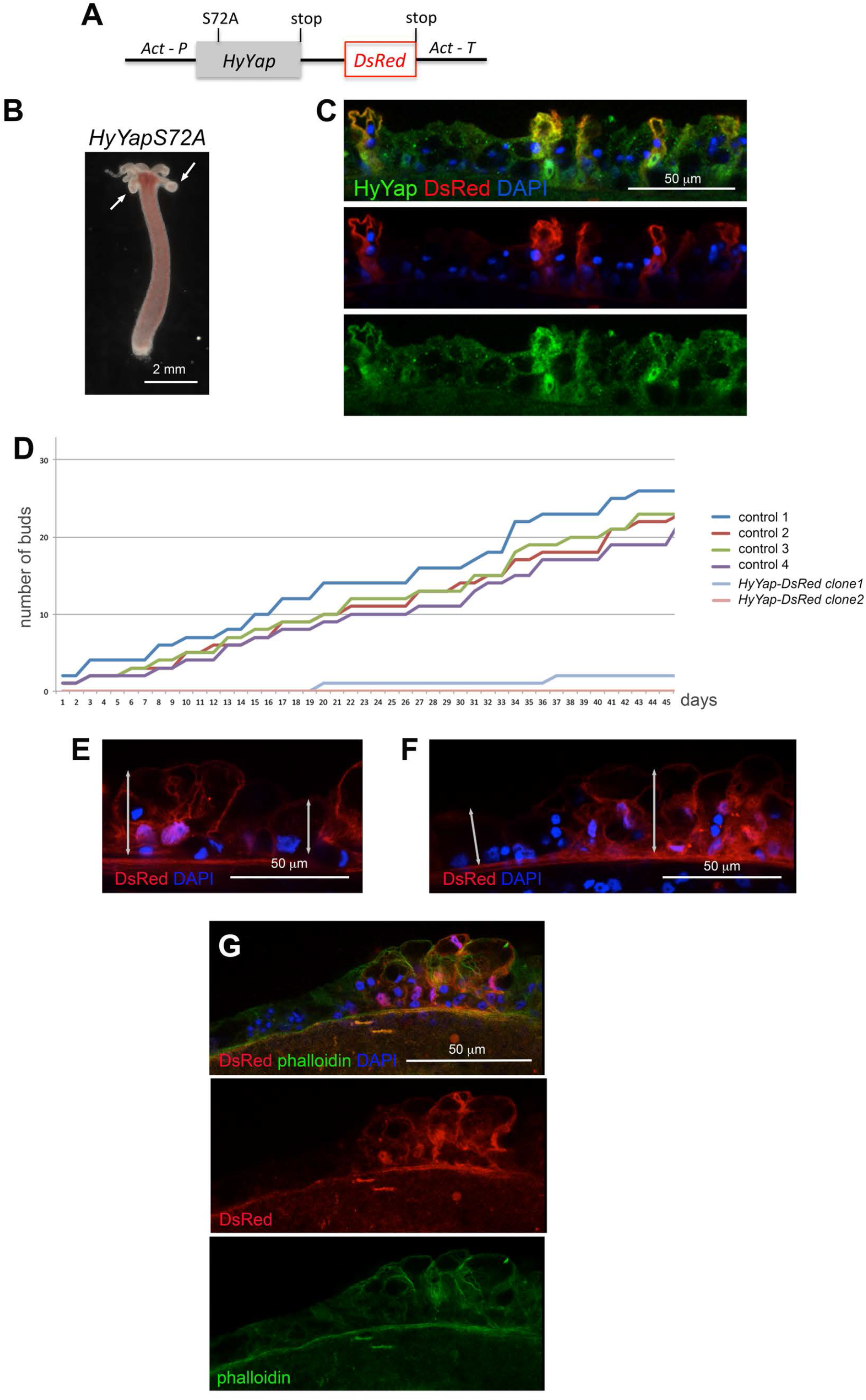
(A) Schematic drawing of the construct used to generate *HyYap-DsRed* transgenic *Hydra*. (B) Photos of live *HyYap-DsRed* hydras. Arrows point to the thick tentacles. (C) Lateral view of the ectoderm of *HyYap-DsRed Hydra* immunostained for HyYap and DsRed. (D) Budding rate of control and *HyYap-DsRed* hydras shown as the number of buds produced by a single polyp over time. (E) Lateral view of the ectoderm of *HyYap-DsRed Hydra* immunostained for DsRed. Double-headed arrows indicate the apico-basal dimension of transgenic and wild type cells. (F) Lateral view of ectoderm of *HyYap-DsRed Hydra* immunostained for DsRed. Double-headed arrows indicated the apico-basal dimension of transgenic and wild type cells. (G) Lateral view of the ectoderm of *HyYap-DsRed Hydra* immunostained for DsRed and phalloidin.

**Figure S4.**
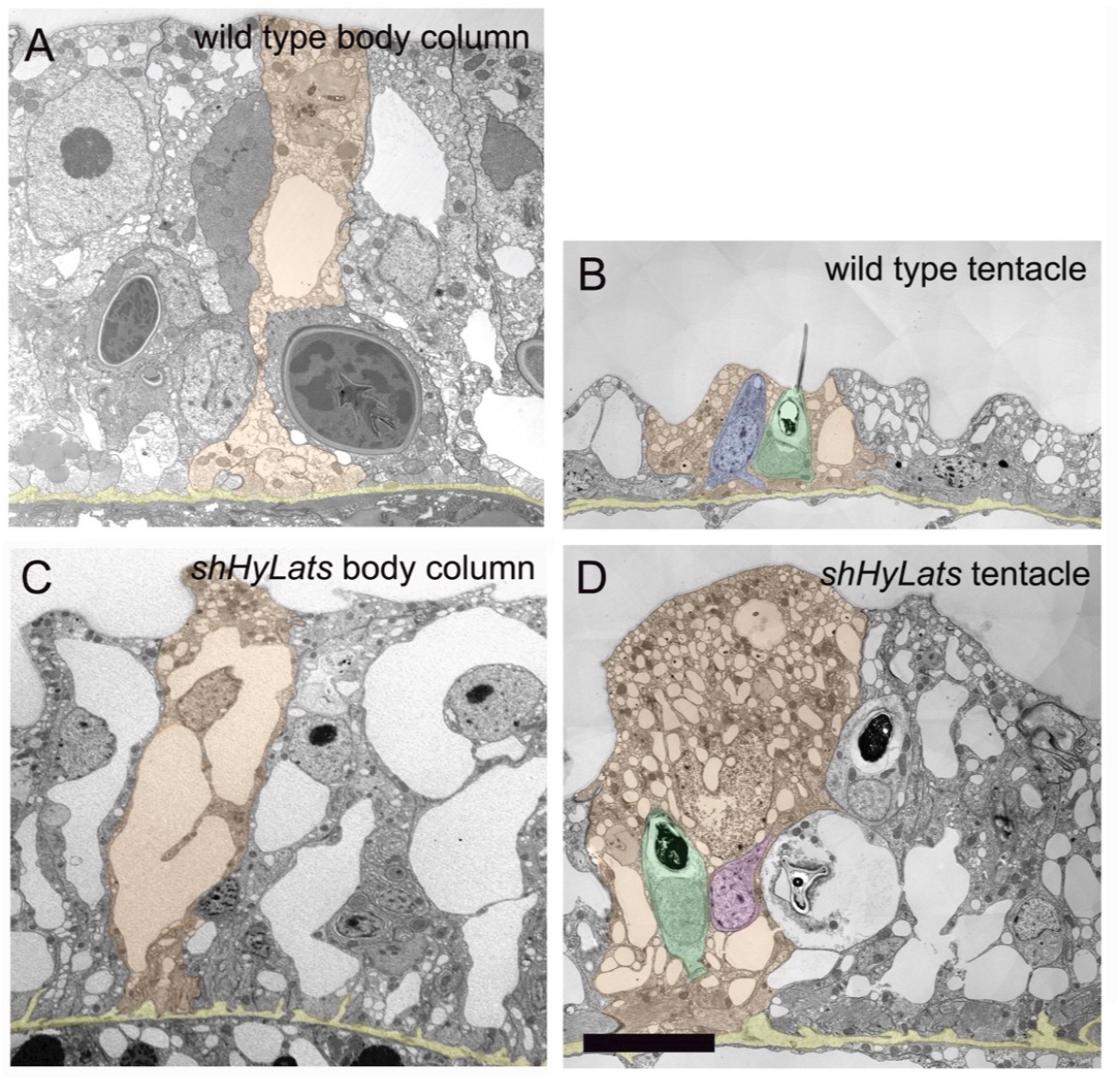
Low magnification transmission electron microscopy images of wild type and *shHyLats* ectoderm. (A,B) Images of body column and tentacle tissue in wild type polyps; (C,D) Corresponding images of *shHyLats* polyps. One representative ectodermal epithelial cell is colored in orange. The mesoglea is colored in yellow. *shHyLATS* thick tentacles show a dramatic expansion of ectodermal epithelial cells in along their apical-basal axis. They generally exhibit a higher volume of intracellular vacuoles indicating that their osmoregulation is affected. (B) Wild type epithelial cells in the tentacles (called battery cells) incorporate differentiated nematocytes (stinging cells; green) mounted at their apical membrane and a sensory nerve cell (blue). (D) Tentacle epithelial cells in *shHyLATS* thick tentacles also exhibit incorporated and fully differentiated nematocytes, but these are usually not mounted at the apical membrane. In addition, these cells show interstitial precursor cells (violet), which are not occurring in the tentacles of wild type polyps. Furthermore, *shHyLATS* polyps exhibit major disruptions of ectodermal basal muscle fibers and mesoglea structure as discussed in detail using high magnification images in the main paper (see Figure 4). Size bar: 20 μm.

## Material and Methods

### Animal and culture condition

The AEP strain of *Hydra vulgaris* was cultured at 18°C in hydra medium (1.0 mM CaCl_2_, 1 mM NaHCO_3_, 0.1 mM MgCl_2_, 0.03 mM KNO_3_, 1 mM TrisHCl pH 7.8, and fed with *Artemia* nauplii every two days.

### *In situ* hybridization

All procedures at room temperature were carried out with rotation on a nutator. Animals starved for 48 hr were relaxed in 2% urethane for 2 min and fixed in 4% paraformaldehyde O/N at 4°C. Samples were washed for 5 min each in 100% ethanol × 3, ethanol:PBT 3:1 × 1, ethanol:PBT 1:1 ×1, ethanol:PBT 1:3 × 1, PBT × 3, following by treatment with proteinase K (10 mg/ml) and 4 mg/ml glycine for 10 min each. Next, smples were treated with 0.1 M triethanolamine (pH 7.8) × 2 and 0.25% (v/v) acetic anhydride in 0.1 M triethanolamine (pH 7.8) × 2 for 5 min each, washed with PBT for 5 min × 2 and postfixed with 4% PFA for 20 min. Fixator was removed by 5 washes with PBT for 5 min. Next, the endogenous alkaline phosphatase was removed by heating samples at 80°C for 30 min and washed sequentially in PBT × 1, PBT:Hybridization buffer (HB) × 1 and HB × 1 for 10 min each. Samples were pre-hybridized in HB at 55°C for 2 hr. Then digoxygenin-labeled RNA probe was added and hybridization was carried out for 48 – 60 hr at 55°C. Hybridization solution (HS) was composed of 50% formamide, 5×SSC (750 mM NaCl, 75 mM sodium citrate), 0.02% (w/v) each Ficoll, bovine serum albumin (BSA, Fraction V), and polyvinylpyrolidone, 200 mg/ml yeast tRNA, 100 mg/ml heparin, 0.1% Tween20, and 0.1% Chaps. To remove unhybridized probe samples were washed at 55°C for 10 min each in HS, HS:2×SSC 3:1, HS:2×SSC 1:1, HS:2×SSC 1:3, following by two 30 min washes with 0.1% CHAPS in 2×SSC. In a preparation for binding with the anti-dioxygenin antibody samples were moved at room temperature and washed twice for 10 min in MAB (100 mM maleic acid, 150 mMNaCl, pH 7.5), then for 30 min in 1% BSA in MAB, and then blocked for 2 hr in blocking solution (BS) (80% MAB-BSA, 20% heat-inactivated sheep serumn). Alkaline phosphatase-conjugated anti-digoxy genin Fab fragments were diluted 1:400 in BS and preabsorbed for at least 2 hr against fixed hydra. The preabsorbed Fab fragments were diluted to a final dilution of 1:2000 in BS and incubated with samples overnight at 4°C. Next day the unbound antibodies were removed by 8 washes with MAB for 30 min each, samples were equilibrated with the alkaline phosphatase staining buffer NTMT (100 mMNaCl, 100 mMTris, pH 9.5, 50 mMMgCl_2_, 0.1% Tween-20) in three 5 min washes, with the final wash also containing 1mM Levamisole. Alkaline phosphatase reaction was carried out at 37°C in NTMT in the presence of 5 ml/ml NBT and 3.75 ml/ml BCIP in the dark. Reaction was stopped with EtOH, refixed in 3.7% formaldehyde, dehydrated in ethanol series 2 min each (70% EtOH × 1, 95% EtOH × 1, 100% EtOH × 2) and mounted in Euparal.

128 – 663 bp segment of Hydra *Wnt3* coding sequence (accession number AF272673) was used to make an *in situ* probe. Digoxygenin-labeled RNA probes were made according to the protocol supplied by Roche.

### Database search and Phylogenetic Analysis

To identify cnidarian homologues of Fat-like, Ds and CELSR proteins we searched the NCBI (http://www.ncbi.nlm.nih.gov) and NHGRI (https://research.nhgri.nih.gov) databases. We have identified *Hydra vulgaris* homologues of Yap (NM_001309649), Hippo (MST) (XM_004212124) and LATS (XM_012698864). Sequences used in the analysis and their accession numbers: DmYorkie (DQ099897), MmYap1 (BC094313), HsYap2 (AAP92710), HsTAZ (AJ299431), NvYap (XM_001627445.2), DrLATS1 (XM_005160312), DrLATS2 (NM_001128256), DmLATS (Warts) (U29608), HsLATS1 (AF104413), NvLATS (XM_001628046), HsMST (U18297), DmMST (Hippo), NvMST (XM_032384310), DrMST (BC164215). For generation of the phylogenetic tree, the sequences were aligned using MAFFT (https://www.ebi.ac.uk/Tools/msa/mafft/) or Clutal Omega (https://www.ebi.ac.uk/Tools/msa/clustalo/) and analyzed using Alaike Information Criterion (http://www.atgc-montpellier.fr).

### Production of antibodies

A peptide corresponding to residues 1 – 159 of HyYap was expressed as GST-fusions using the GEX4t-1 vector (Millipore) in *E. coli* strain BL21 and purified on glutathione-agarose (Thermo Scientific). Purified protein was used to immunize guinea pigs (Cocalico Biologicals).

### Immunoblot and Immunofluorescence analysis

For immunoblotting, polyps were dissolved in lysis buffer (5% SDS; 10% glycerol; 60 mM Tris-HCl, pH 6.8) containing 2% beta-mercaptoethanol, boiled for 5 min, chilled on ice for 5 min and electrophoresed in a 10% SDS-PAGE polyacrylamide gel. Transfer of the proteins onto a PVDF membrane was done in transfer buffer (20% methanol, 25 mM Tris base, 192 mM glycine) overnight at 4° C. The membranes were incubated with antibodies (total anti-HyYap serum was used at 1:1000 dilution, anti-actin (clone C4, Millipore) at 1:2000 dilution, anti-GAPDH (Sigma) at 1:1000 dilution) in blocking buffer (TBS-tween 0.1% containing 5% powdered milk) for 90 minutes at RT. After 3×10 min washes with TBS-T, membranes were incubated with HRP-conjugated secondary antibody (GE Healthcare) diluted 1:10000 in blocking buffer for 1 h at RT. Visualization was done by ECL detection (Thermo Scientific).

To perform immunocytochemistry on whole mounts animals were relaxed in 2% urethane for 2 minutes, and then fixed in either Lavdovski’s fixative (ethanol:formaldehyde:acetic acid:H_2_O 50:10:4:36) for HyYap and GFP antibodies, or in 4% paraformaldehyde for phalloidin staining overnight at 4°C. Then, animals were washed 3 × 10 min in PBT (PBS with 0.1% Triton), animals fixed with 4% paraformaldehyde were permeabilized in PBS with 1.0% Triton for 30 min, and incubated in PBT with 2% BSA for 1 hour. Samples were incubated with primary antibodies overnight at 4°C. Then, animals were washed 3 × 10 min PBT, incubated with secondary antibodies for 30 min, washed with PBT 3 × 10 min and mounted using Vectashield mounting medium containing DAPI (Vector Laboratories, Inc). Primary antibodies dilutions were as follows: anti-HyYap total serum, 1:1000; anti-GFP (Abcam, ab13970). Fluor-conjugated secondary antibodies (Jackson Laboratory) were used at 1:400 dilution. Alexa555-phalloidin (Abcam) was used at 1:2000 dilution.

### Measurement of immunofluorescent intensity

To measure the intensity of immunofluorescence 1.5 mm z-stack confocal images were analyzed by NIS-Elements AR Analysis.

### *shRNA* production and electroporation

*shRNAs* were designed and made according to [1]. For each gene two hairpins were synthesized (See Table S1) and both were used for electroporation at 1:1 ratio. Electroporation procedure was performed according to [2].

